# Reorganization of bird communities along a rainforest elevation gradient during a strong El Niño event in Papua New Guinea

**DOI:** 10.1101/2022.07.26.501620

**Authors:** Katerina Sam, Leonardo Ré Jorge, Bonny Koane, Richard Hazell, Phil Shearman, Vojtech Novotny

**Author notes:** Corresponding author: Katerina Sam –, Biology Centre AS CR, v. v. i., Institute of Entomology, Ceske Budejovice 370 05, Branisovska 31, Czech Republic. Author Contributions: KS – conceived and designed the experiment, KS conducted the surveys and wrote the manuscript. BK – conducted the surveys. PS – provided climatological data. RH – conducted the part of the survey after El Niño. LRJ conducted statistical analyses and wrote the manuscript with KS. VN– supervised initial project of KS, financially supported the work, provided logistical support. All authors commented on the manuscript.

## Abstract

The El Niño 2015 event, most extreme since 1997, led to severe droughts in tropical wet Papua New Guinea (PNG), reducing May to October dry season rainfall by - 75% in the lowlands and 25% in the highlands. Such droughts are likely to have significant effects on terrestrial ecosystems, but they have been poorly explored in Papua New Guinea. Here we report changes in bird community composition prior, during and after 2015 El Niño event along the elevational gradient ranging from 200 m to 2,700 m a.s.l. at the Mt. Wilhelm rainforest in PNG. The abundance of birds in lowlands dropped by 60% but increased by 40% at elevations above 1700m during El Niño year. In the following year, the individual bird species reached mean population sizes similar to pre-El Niño years but did not fully recover. Species richness roughly followed the pattern of observed abundance and quickly and fully re-established after the event to the pre-El Niño values. Thus, at least some terrestrial birds seem to react quickly to the extreme droughts in lowlands and shifted to less affected mountain habitats. We recorded upper elevational range limits to shifts by more than 500m asl in 22 bird species (out of 237 recorded in total) during El Niño year, in contrast to their typical ranges. Our study suggests that a strong El Niño event can have strong but reversible effects on bird communities as long as they have an opportunity to move to more favourable sites through undisturbed habitats.

## Introduction

El Niño-Southern Oscillation (ENSO) is an example of a large-scale climate change that occurs repeatedly, in average once every five years, with varying effects in different places in the world. The ENSO cycle, including both El Niño and La Niña, causes global changes in temperature and the redistribution of rainfall. Northern and central South America becomes predominantly dryer during El Niño evens. Its global effects further include also drier than normal weather from northern Australia to SE Asia. In contrast, rainfall is generally restricted to the coast, west of the Andes, where it raises above normal (Holmgren et al. 2001). In terms of temperature, El Niño is associated with a band of warm ocean water that develops in the central and east-central equatorial Pacific (Cai et al. 2019). The opposite of El Niño, La Niña, occurs when sea surface temperatures in the central Pacific drop to lower-than-normal levels.

Specifically, El Niño is associated with high rainfall in South America led to increases in abundance and increasing population trends of vertebrates such as small mammals and carnivorous birds that prey on them (Lima et al. 1999a, Lima et al. 1999b, Jaksic et al. 1997, Meserve et al. 1999), or other land birds (Jaksic and Lazo 1999, Wolfe et al. 2015, Jones et al. 2019). However, several studies reported negative relationship and decline of bird abundances (Blake and Loiselle 2015, Santillán et al. 2018) or their annual survival (Ryder and Sillett 2016) in a period of increased rainfall. In Asia and Australia, El Niño events bring warm, dry, sunny conditions to a large portion of the usually wet tropics. Similarly to South America, also here the unusually dry and sunny conditions may have several different effects on food resources which can influence various taxa (including birds) for months or even years (Appanah 1985, 1993, Owens 1995, Wright et al. 1999, Butt et al. 2015), but the phenomenon is there less studied.

As noted above, ENSO associated long- and short-term fluctuations in rainfall can affect terrestrial endotherms in multiple ways. Both via direct (physiological) and indirect (biotical) mechanisms underlying the rainfall-demographic interactions (Boyle et al. 2020). Interpretations of the biotic mechanisms are pervaded by the idea that food availability drives dynamics of bird populations. For example, interannual variation in precipitation affect arthropods directly, as too much rain reduces activity of arthropods, making them less available to birds to consume. Further, precipitation affects higher trophic levels through production of leaves, flower, fruits, and seeds, which in turn provide rich source of food for arthropods (Butt et al. 2015, Immelmann 1969). Despite production of fruits and other plant tissues is more directly linked to water availability (Sakai et al. 2006, Gandiwa et al. 2016), and the abundances of their consumers, e.g. arthropods, are mediated by them, it is hypothesised that birds feeding directly on plant products could be more quickly and directly affected than insectivorous birds (Kishimoto-Yamada et al. 2015). However, too much rain can lead to direct foraging limitation of birds, irrespective of whether a species is an insectivore or frugivore.

Dietary shifts during extended droughts are poorly reported for birds, but it is likely that some occur (Sam et al. 2017, Tebbich et al. 2004). In general, droughts and extended dry periods reduce bird diversity and abundance (Herremans 2004), either through opportunistic habitat shift or migration (Dean 2004) or lowered survival (Hanmer 1997). Frugivorous birds can be then in advantage as they are evolutionarily predisposed for migrations following phenology of fruiting trees, despite some frugivores at low elevations do not move (Levey and Stiles 1992). On the other hand, tropical insectivores are believed to be more sedentary than frugivores (Sekercioglu et al. 2002), as abundances of arthropods rarely become a limiting factor. We can thus expect different mechanism being linked to precipitation and demography of birds belonging to different feeding guilds or even species (Boyle et al. 2020).

Further, the difference in rainfall regime can have different effects in more seasonal and aseasonal regions or habitats. In arid tropical or more seasonal tropical regions, food abundance tightly underlies demographic responses to interannual variation in rainfall (Gandiwa et al. 2016) and bird communities are adapted to such fluctuations, being potentially less affected by sudden changes in climatic conditions. In far wetter tropical regions, precipitation paramount in shaping biotic seasonality. Low rainfall leads to low plant resources which cascade to reduced populations in higher trophic levels, from producers to birds (Reich 1995, Wright et al. 1999, Stouffer et al. 2013). Alternatively, low rainfall is also sometimes associated with lowered plant defence and subsequent insect outbreaks during rains following the droughts potentially affecting higher trophic levels positively (Van Bael et al. 2004). There, animals used to relatively stable conditions, e.g. in lowlands for tropical elevational gradient, might be eventually more negatively affected by unusual climatic conditions associated with unpredictable El Niño events (Jaksic and Feinsinger 1998, Styrsky and Brawn 2011, Boyle et al. 2020) than at higher elevations. This might be because the birds at higher elevations are used to more unstable conditions naturally, or because the high elevations are rarely severely affected by El Niño related droughts (Cai et al. 2019).

Although tropical humid mountains, similar to Mt. Wilhelm, are usually not limited by water availability, rain forests on Mount Kinabalu (Borneo) were seriously affected by the 1997–98 El Niño drought (Aiba and Kitayama 2002), even though it was not as severe as the 2015 El Niño (WFP 2015). Such changes in rainfall distribution significantly affect avian reproduction and survival, fecundity and other life-history traits (Sillett et al. 2000, Mendoza et al. 2017).

In Papua New Guinea (PNG), the El Niño started developing during the last months of 2014 as a weak event, continued to strengthen after Typhoon Higos in February 2015, and peaked in August 2015 (ENSO 2015). Specifically, the last months of 2014 and early months of 2015 brought lower than usual rainfall, and the wet season ended early. The 2015 El Niño event was considered to be “very strong” (i.e. 2.0°C or higher sea surface temperature anomaly; Null 2017). It was the strongest El Nino event in PNG since the event in 1997. During May 2016, the El Niño event vanished as near average to below average sea surface temperatures expanded across the eastern equatorial Pacific Ocean (IOM 2015, ENSO 2016). PNG has a rugged topography, which might lead to different effects of El Niño in different areas. In this sense, contrasting effects of climate extremes at high and low elevations were subsequently reported on plant phenology (Null 2017). During the dry season (May-to-October) in 2015, rainfall was severely below normal in the lowlands (where rainfall was only 25% of the long-term average) but closer to normal in highlands (rainfall ca. 75% of long-term average). Additionally, the highest elevations of PNG also suffered from unusual local frosts. The whole elevational gradient of Mt. Wilhelm, in the Central Range, spans from the alluvial forest (<500 m a.s.l.), through wet foothill forest (ca. 500–1,500 m a.s.l.), lower montane forest (ca. 1,500–2,500 m a.s.l.), and up to upper montane forest (ca. 2,500-3,500 m a.s.l.) and all the way to tree line which is at 3,700 m a.s.l.. Typically, precipitation at the sites is slightly seasonal, with a mild dry season from May till October (154 ± 48 mm of rainfall per month) and a wet season from November to April (515 ± 158 mm of rainfall per month). The lowland and mid-elevation forests up to 1,700 m are usually more seasonal, while higher elevations have a more stable rainfall regime (221 ± 65 mm in “dry season” and 338 ± 98 mm in “wet season”) due to the condensation zone, and nearly daily afternoon rains throughout the year.

A major problem with El Niño studies is that they often fail to cover the entire period before, during and after El Niño due to their unpredictable and relatively rare occurrence (but see e.g. (Wright et al. 1999, Jaksic 2001). The study of after El Niño events is particularly important, as they can inform us about the resilience of animals and plants, and their ability to react flexibly to climate changes. Before El Niño events are important for setting a baseline standard of comparisons. However, it is difficult to predict when the El Niño will start and thus when the sampling should be started effectively. El Niño events also vary markedly in strength, which further complicates the assessment of their importance for ecosystem dynamics. The 2015 event provided the best opportunity to study a strong El Niño event in the past 20 years.

Here, we report on the effect of this extreme El Niño event on forest bird communities along an elevational gradient in Papua New Guinea. Specifically, we describe the communities of birds belonging to four different guilds and show the changes to their elevational distribution before, during and after an El Niño event along a tropical rainforest gradient, as we expect different indirect effect of the even on different deeding guilds. We hypothesize that bird response to El Niño could be of 3 types: indifferent, if bird distribution along the gradient did not vary along our sampling; resilient, if bird distribution changed during the El Niño event, and then returned to the previous state; and long-term impacted, if the distribution of the birds was different respectively before, during and after the El Niño event. We expected that some species will decline in abundances for long time and have lower survival during strong El Niño event, while only some species may be resilient, as there are many studies showing that tropical birds are typically not able to rapidly shift their distribution to track climatic niche (i.e., suitable humidity and temperature; Chen et al. 2011, Freeman and Freeman 2014). Additionally, we tested whether birds belonging to different guilds, and thus dependent on different types of resources, had different elevational distributions and responses to the El Niño event.

## Methods

Our study was performed on the slopes of Mt. Wilhelm (4,509 m a.s.l.) in the Central Range of Papua New Guinea. The whole transect is 20 km long and consists of eight sites, evenly spaced at 500 m elevational increments from the lowland floodplains of the Ramu river at 200 m a.s.l. to the timberline at 3,700 m a.s.l. In this study, we surveyed lower six elevations between 200 and 2700 m a.s.l. only. (Fig. 1). Within the surveyed transect, the habitats are tropical lowland alluvial forest (200 m a.s.l.), tropical foothill forest (700 and 1,200 m a.s.l.), lower montane moss forest (1,700 – 2,200 m a.s.l.) and upper montane forest (2,700m a.s.l.) according to (Paijmans 1976). The typical species composition of the forest (Paijmans 1976), general climatic conditions (McAlpine et al. 1983) and individual study sites (Sam et al. 2014) are described elsewhere. Along the gradient, density of shrubby understory and canopy openness increases with increasing elevation (Fig. S1). Average annual precipitation is 3,761 mm in the lowlands (200 m, Usino Junction meteorological station at - 5.56283, 145.3542), rising up to 4,300 mm at 2,700 m a.s.l., with a distinct condensation zone between 2,200 and 2,700 m a.s.l (Mondia Pass meteorological station at 5° 49’ 34.729”E 145° 8’ 1.388”) during non-El Niño years (Fig. 2). During the 2015 El Niño, the largest fluctuations in total annual rainfall were observed at 200 and 2,700m, where it dropped significantly to 2,122 mm and 2,503 mm from typical 3,761 mm Fig. 2). A decrease in the amount of rainfall was detectable at 200 m already in 2014 (Fig. 2). The El Niño in 2015 and 2016 caused widespread severe droughts (−75% of rainfall) in lowlands and foot-hills, mild droughts in lower montane forests (−25% of rainfall) and unusual frosts in upper montane forests (above 3,000 m a.s.l., i.e. not surveyed in this study) of PNG, where the rainfall was however not significantly affected.

**Figure 1.**
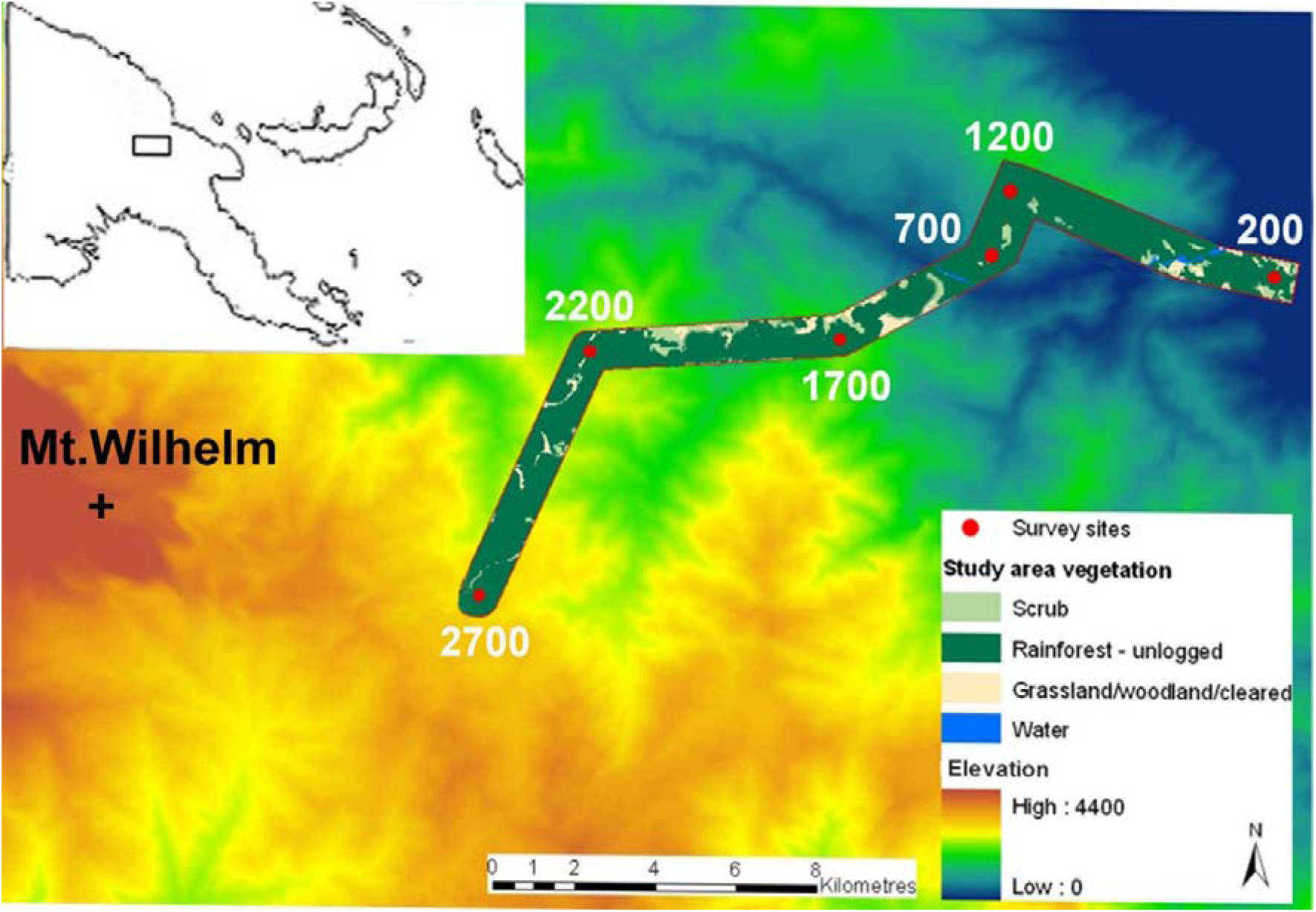
Map of the study sites. The study sites were selected to represent the typical forest habitats of Mt Wilhelm. The sampled rain forest gradient spanned from the lowland floodplains of the Ramu river (Kausi, 200 m a.s.l., 5° 44’ S 145° 20’ E) to the 2700 m a.s.l. (Bruno Sawmill, 2700 m a.s.l., 5° 48’ S 145° 09’ S). Source of the data for the map: (Shearman et al. 2009, Bryan et al. 2015)

**Figure 2.**
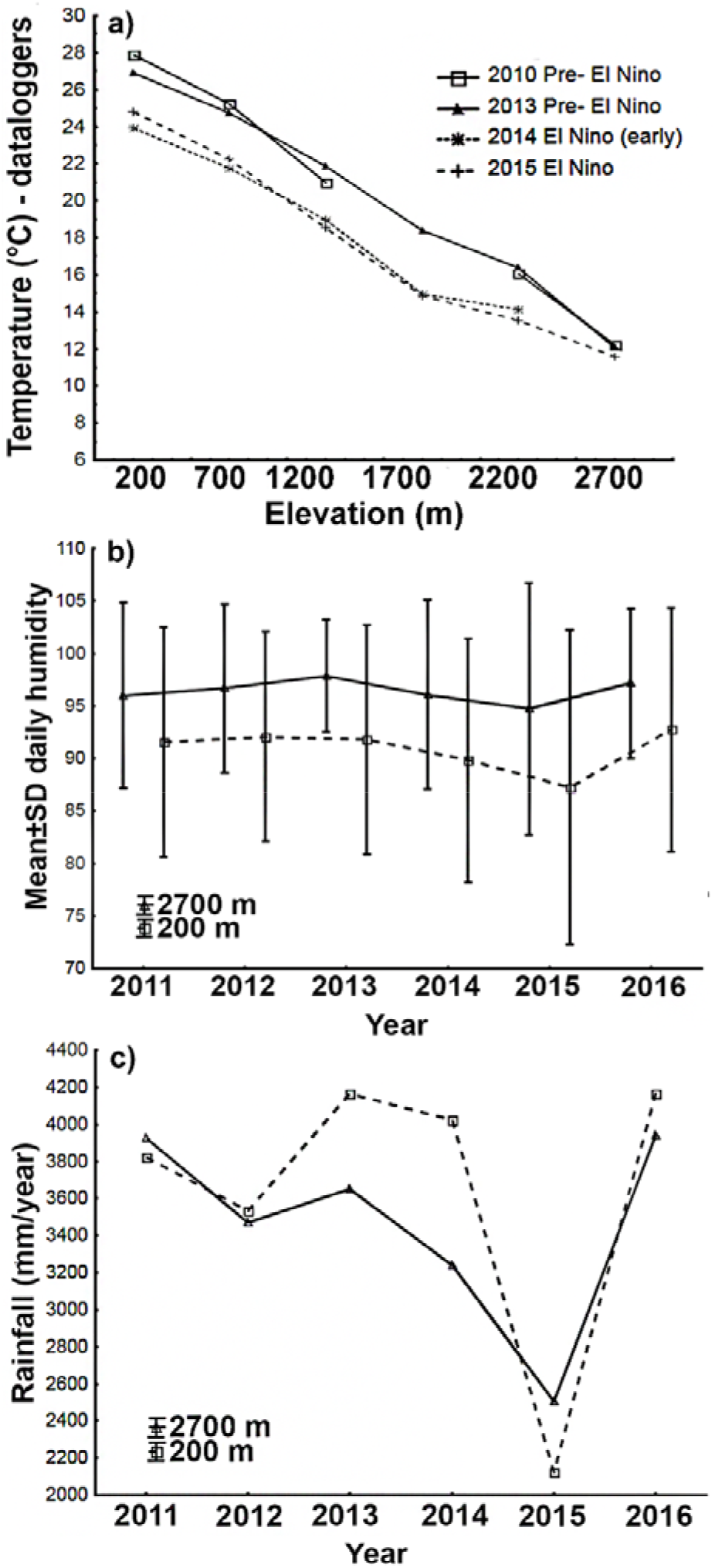
Mean daily temperatures for forest interior of Mt. Wilhelm gradient study sites **(a)** and mean (±s.e.) daily humidity **(b)** at two study sites across several years (measured by dataloggers Comet R3120). Mean rainfall **(c)** in mm per year at 200 and 2,700 m a.s.l.; data from Usino Junction and Mundiapass meteorological station provided by Phil Shearman - (Bryan et al. 2015).

Mean annual temperature and humidity were measured by Comet R3120. Temperature decreases usually from 27.4°C at the lowland site to 8.37°C at the tree line at a constant rate of 0.54°C per 100 elevational metres (Fig. 2) during non-El Niño years. During the El Niño in late 2014 and in 2015, temperature was on average 3 °C lower than in before El Niño years (2010 and 2013) at four lower elevations of the gradient (Fig. 2). This change was driven by relatively colder nights (24.5 vs. 21.3°C), while days remained similarly hot (27.8 vs. 26.8) during before El Niño (2010) and El Niño (2015). At higher elevations (2,200 and 2,700 m), the difference between the before El Niño and El Niño years was 2.4 and 0.96 °C respectively (Fig. 2). Unfortunately, data from higher elevations are not available for El Niño years.

### Bird sampling

We surveyed bird communities using point counts. We carried out the bird counts at each elevational site at 13 points regularly spaced along a 2,050 m transect (successive points were 150 ± 5 m apart to avoid overlap). We led the transects along newly prepared trails, with maximum elevational extent ±60 m elevation (Sam et al. 2019). We surveyed birds only during days without rain. We recorded all birds seen or heard within a fixed radial distance of 0 - 50 m (estimated or measured by a laser rangefinder). We started censuses 15 min before sunrise and each count lasted 15 min so that all 13 points were surveyed before 10:30. All three observers (KS, BK, RH) had between 3 to 8 years of previous experience working with birds in Papua New Guinea (Sam and Koane 2014, Sam et al. 2014, Peck et al. 2017, Sam et al. 2019). K.Sam and B. Koane surveyed birds together in 2010-2015, R. Hazell joined them in 2015 and 2016. We also recorded vocalizations at each survey point during surveys using a Marantz PMD 620 recorder (Eindhoven, Holland) and a Sennheiser ME67 microphone (Hanover, Germany). This allowed us to identify any bird vocalizations unrecognized during the point count surveys. We followed the IOC World Bird List (version 6.1.; www.worldbirdnames.org) species-level taxonomy and nomenclature.

We conducted four surveys during the four years of study. During each survey, we conducted point-counts at 13 points per elevation and replicated these counts thrice in three consecutive days. The before El Niño surveys were conducted between 15th September and 15th October of 2010 (first three consecutive days from 2010) and 15th September and 15th October of 2013 (first three consecutive days from a larger study in 2013) by Sam et al. (2019). The third survey was conducted during the El Niño event, between October 1^st^ 2015 and December 2^nd^ 2015. The last, after El Niño survey was conducted between 8^th^ of April and 6^th^ of June 2016, thus aiming for the typical end of wet season which usually weakens in April/May. Thus, the one-month long surveys in 2010 and 2013 were conducted in the same month and the survey in 2015 started just 15 days later. The analyses are based on 13 data points (i.e., physical locations) per survey year at each elevational study site, each of the data points representing a pooling of the three consecutive point-counts. The summed abundances closely correlated with maximal abundances sometimes used by other authors in similar studies (Fig. S2). Unfortunately, the survey in 2016 was conducted in different months than previous surveys. The timing was given by the funding body supporting the logistics of post-El Niño survey. While interpreting the results, it should be kept in mind that this survey thus isn’t fully comparable with the rest of the survey. However, according to our previous observations, bird activity and behaviour is similar at the beginning and at the end of the wet season, when we held our surveys.

In all analyses, we used the total number of species with standardized area and time from all three surveys. The observed species richness is a good index of actual species richness, as long as it is based on sufficient and standardized sampling effort (Gotelli and Colwell 2001). However, species richness indices can capture different aspects of diversity and thus produce different results compared to standardized sampling effort (Gotelli and Colwell 2001). However, considering the potential issues with the detectability of birds and given that species richness is often correlated with sampling effort, we obtained Chao 1 estimates to confirm that use of observed species richness doesn’t underestimate undetected species significantly, as recommended by Gomez et al. (2019). We used EstimateS (Colwell, 2009) program to obtain Chao 1 estimates and its confidence intervals by randomizing the count matrix 1,000 times. Then we concluded that estimated species richness corresponds very closely with the raw number of detected species detected (Fig. S3), which are safe to use in analyses.

Recorded birds were partitioned into four trophic guilds: insectivores, frugivores, frugivore-insectivores (e.g., catbirds which feed occasionally on fruits and insect), and nectarivore-insectivores (e.g., honeyeaters which feed on nectar and small insect, and which included also pure nectarivores) based on published dietary information (Table S1; Sam et al. 2017, Sam et al. 2019). We used four guilds to be able to detect fine differences between strategies, which would be impossible if all mixed feeders are considered as omnivores as it is done in several other studies (e.g., Adeney et al. 2006). Only forest species, i.e., those actively using the forest habitat but not those flying across, were included in the analyses. All raptors and swifts were excluded (113 individuals of 10 species, Table S1) since it was difficult to sample them in a standardized manner from the forest interior (Sam et al. 2019).

#### Data analysis

We used generalized linear mixed models (GLMM) to describe the effects of elevation and presence of El Niño events on the abundance and richness of birds in our surveyed plots. We held all analyses for the overall richness and abundance of birds, and also separately for each of the four guilds considered. As our richness and abundance data has no zeroes for the overall analysis and none or a very small proportion of zeroes for all the guild specific data, we did not consider zero-inflated models.

Each of the response variables (either richness or abundance) was modelled with a Poisson error distribution and log link function. The models for abundance had a moderate level of overdispersion based on what we decided for Poisson distribution. Further, negative binomial models fitted to the data showed the exact same ranking of models and very similar performance overall. Thus, we used Poisson mixed models for all analyses.

We used a model selection approach to test for the different hypotheses described in the introduction. We built a set of candidate models including the effect of elevation and/or the El Niño event; 1) Elevation was modelled either as a linear or a second order orthogonal polynomial, to test whether the bird distributions were monotonic or hump-shaped along the elevation gradient. 2) To consider the three possible responses of the bird community to the El Niño event, we used two different codings of the sampling years. The rationale here is that, instead of looking at post hoc comparisons between years to validate our hypotheses, we are testing them directly through different models. Thus, if birds are a) indifferent to the El Niño event, none of the models including any coding of temporal variation will have the best performance. If birds are b) resilient to El Niño, a simpler coding of temporal variation, considering only whether the sampling was done during El Niño or not, will have a better performance than a variable including both before, during and after El Niño as different levels. If birds have c) long-term effects, the more complex temporal variable with before, during and after levels will perform better than models with no effect or the more simple effect of El Niño.

We considered all the model combinations of Elevation and El Niño effects alone, with an additive effect and also with an interaction term. We never included both El Niño effect variables together, but rather as two competing explanations for the effect of El Niño. This yielded a total of 13 candidate models for each response variable (Table 2). For all models, including the null, we included the year/elevation combination as a random factor, allowing the 13-point count locations to be treated as a proper replicate, with the random factor accounting for possible correlations within transects. It was not possible to add year by itself as a random factor, given the low number of years that prevents a correct estimation of the variance term.

We employed the corrected Akaike Information Criterion (AICc; Burnham & Anderson 2002) on the same set of candidate models for all response variables. After fitting, the models were tested for collinearity of the predictors using the performance package (Lüdecke et al. 2011). All analyses were held in the R environment version 4.0.3, using the package lme4 (Bates et al. 2014) to build the models and bbmle (Bolker & Bolker 2012) to calculate AICc (Burnham and Anderson 2002).

## Results

We documented 17,149 bird individuals from 237 species (Table S1). The shape of abundance distribution along the gradient changed from a monotonic decline in abundance with elevation both prior and after El Niño to a strongly hump-shaped gradient with a peak at mid-elevation (Fig. 3). The model testing long term effect of El Niño had better explanatory power than the model considering resilience of abundances of birds (Table 1). In contrast, species richness seemed to resilient to the effect of El Niño (Table 1).

**Figure 3.**
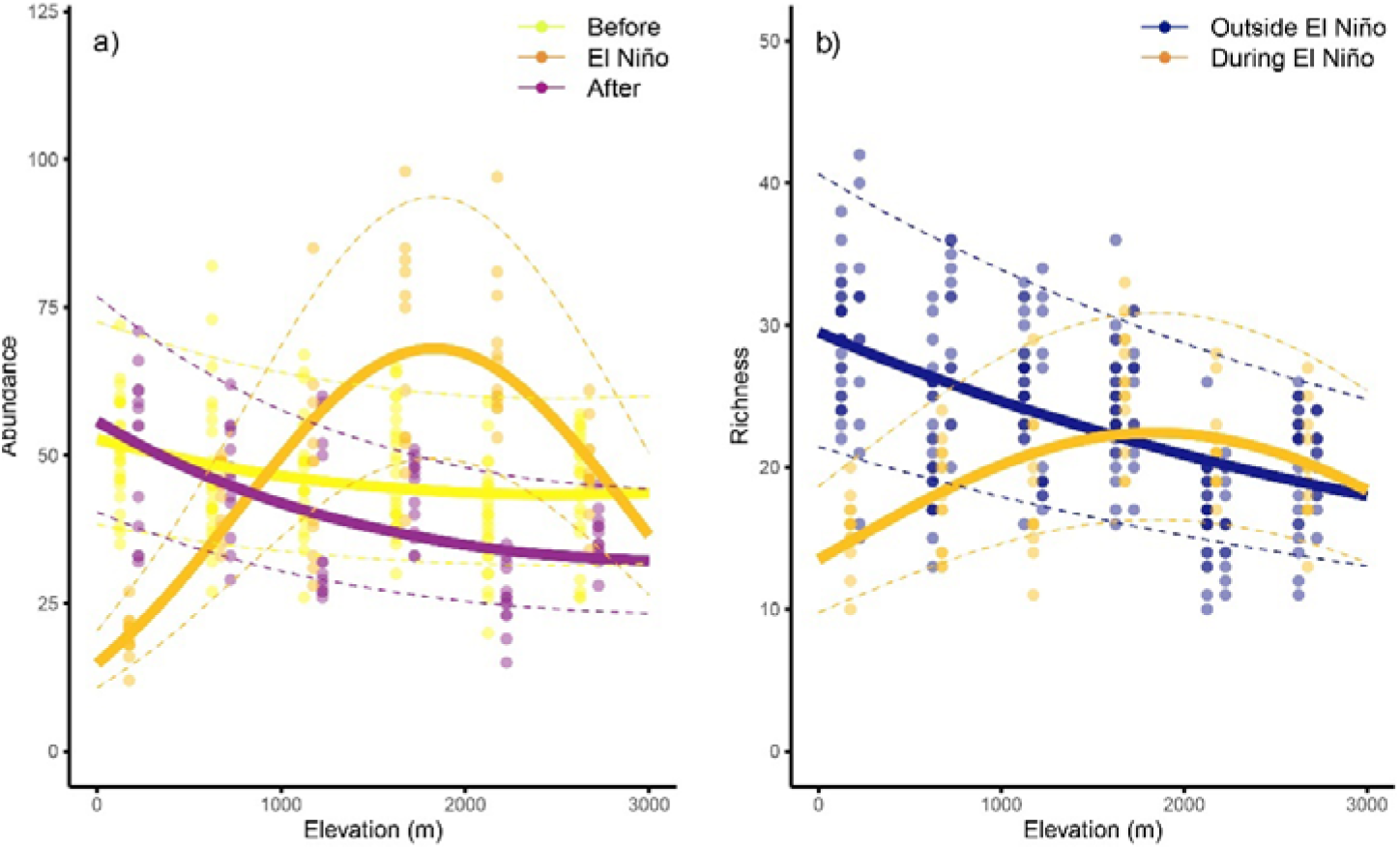
Abundance **(a)** and species richness **(b)** of birds recorded in each time period (colour coded) at 13 points spatially distributed at each of the elevational study sites. Abundances and species richness are expressed as number of individuals recorded in 45 minutes per 0.78 ha, i.e. pooled from 3 surveys per point with radius 50 m. Each of the points was surveyed for 15 minutes per day and the survey at each point was replicated three times in three consecutive days every year. Fitted lines represent the fit from the best model from Table 1 and dashed lines represent 95% confidence intervals of the predicted models.

**Table 1.** Candidate models affecting abundance and richness of birds along an elevational gradient in Papua New Guinea. All models are generalized linear mixed models and include all the year/elevation combinations as a random intercept factor in addition to the fixed factors listed on the table. The null model includes only a fixed intercept in addition to the random factor. Factors and their combinations having effect on observed species richness and abundance. Effect of elevation is considered as polynomial or linear. *Long-term* effect of El Niño in models is represented by three levels – prior to, during, and after El Niño, while *resilience* is represented by 2 levels only – outside and during El Niño. Confidence intervals for effects of the selected models are noted in Table

|  | Abundance |  |  | Species richness |  |  |
| --- | --- | --- | --- | --- | --- | --- |
|  | dAICc | df | weights | dAICc | df | weights |
| elevation<sup>2</sup> * resilience | 4.1 | 7 | 0.114052 | 1.1 | 7 | 0.05127 |
| elevation<sup>2</sup> * long-term | <u>0</u> | <u>10</u> | <u>0.885946</u> | 6.9 | 10 | 0.00269 |
| elevation * resilience | 19.9 | 5 | <0.0001 | <u>0</u> | <u>5</u> | <u>0.93178</u> |
| elevation * long-term | 20 | 7 | <0.0001 | 3.7 | 7 | 0.00568 |
| elevation | 29.7 | 3 | <0.0001 | 10.8 | 3 | 0.00198 |
| elevation + resilience | 31.5 | 4 | <0.0001 | 9.4 | 4 | 0.00219 |
| elevation<sup>2</sup> | 30.7 | 4 | <0.0001 | 12.6 | 4 | 0.00079 |
| elevation<sup>2</sup> + resilience | 32.6 | 5 | <0.0001 | 11.2 | 5 | 0.00091 |
| elevation + long-term | 32.1 | 5 | <0.0001 | 11.4 | 5 | 0.00091 |
| elevation<sup>2</sup> + long-term | 13.2 | 6 | <0.0001 | 33.1 | 6 | 0.00003 |
| resilience | 12.6 | 3 | <0.0001 | 29.5 | 3 | 0.00005 |
| long-term | 30 | 4 | <0.0001 | 14.5 | 4 | 0.00059 |
| null | 29.6 | 2 | <0.0001 | 13.4 | 2 | 0.00089 |

During El Niño, the abundances (median±s.e. across 13 points of the given elevational site) of birds from 200 m a.s.l. experienced a large drop by ca. 60% in their abundances to median of 19.61±0.98 from 51.61±1.77 of their before El Niño abundance, calculated as median of 2010 and 2013 (Fig. 3). After El Niño, their median abundances there were however similar to those before El Niño (51.15±3.76, Fig. 3). In contrast, bird abundances at 1700 and 2200 m increased by ca. 40% during El Niño to 66.66±3.35 and 71.38±6.58 from 47.42±2.01 and 39.15±1.75 typical abundances before El Niño respectively (Fig. 3). In the following year, the individual bird species reached mean population sizes nearly identical to prior El Niño years. It is however notable that the abundances along the elevational gradient in the years before the El Niño event were overall ca. 30% higher compared to the after El Niño year (Fig. 3).

As expected, the species richness gradients roughly followed the patterns observed for abundance, with a very strong mid-elevation peak during the El Niño, and a monotonic decline of diversity from the lowland maximum along the gradient both before and after the El Niño event (Fig. 3b). Resilience of the species richness was supported by the models, as shown by the better performance of the model including simple presence and absence of El Niño over the model separating also before and after El Niño years. This indicates the species richness pattern returned quickly to their original state (Table 1).

Abundances (Fig. 4) and species richness (Fig. 5) of individual feeding guilds were also influenced by El Niño, except nectarivores and nectarivore-insectivores (Table 2). However, it is important to note that there is top model only for abundances of frugivores and insectivores. For species richness and abundances of other feeding guilds, the resilient and long-term models have equal explanatory power (Table 2). Within individual feeding guilds, there is thus no clear support for either the resilience or long-term models. In case of frugivores and frugivore-insectivores, the abundance, and species richness outside of El Niño decreased monotonically and showed a hump-shape during El Niño (Fig. 4, Fig. 5). In case of insectivorous birds, the abundance and species richness showed a hump-shaped patter in all years, but its peak moved to higher elevations during the El Niño year (Fig. 4, Fig. 5). The insectivorous birds showed generally the lowest fluctuations between the years, and the observed patterns were driven by fruit-feeding guilds.

**Figure 4.**
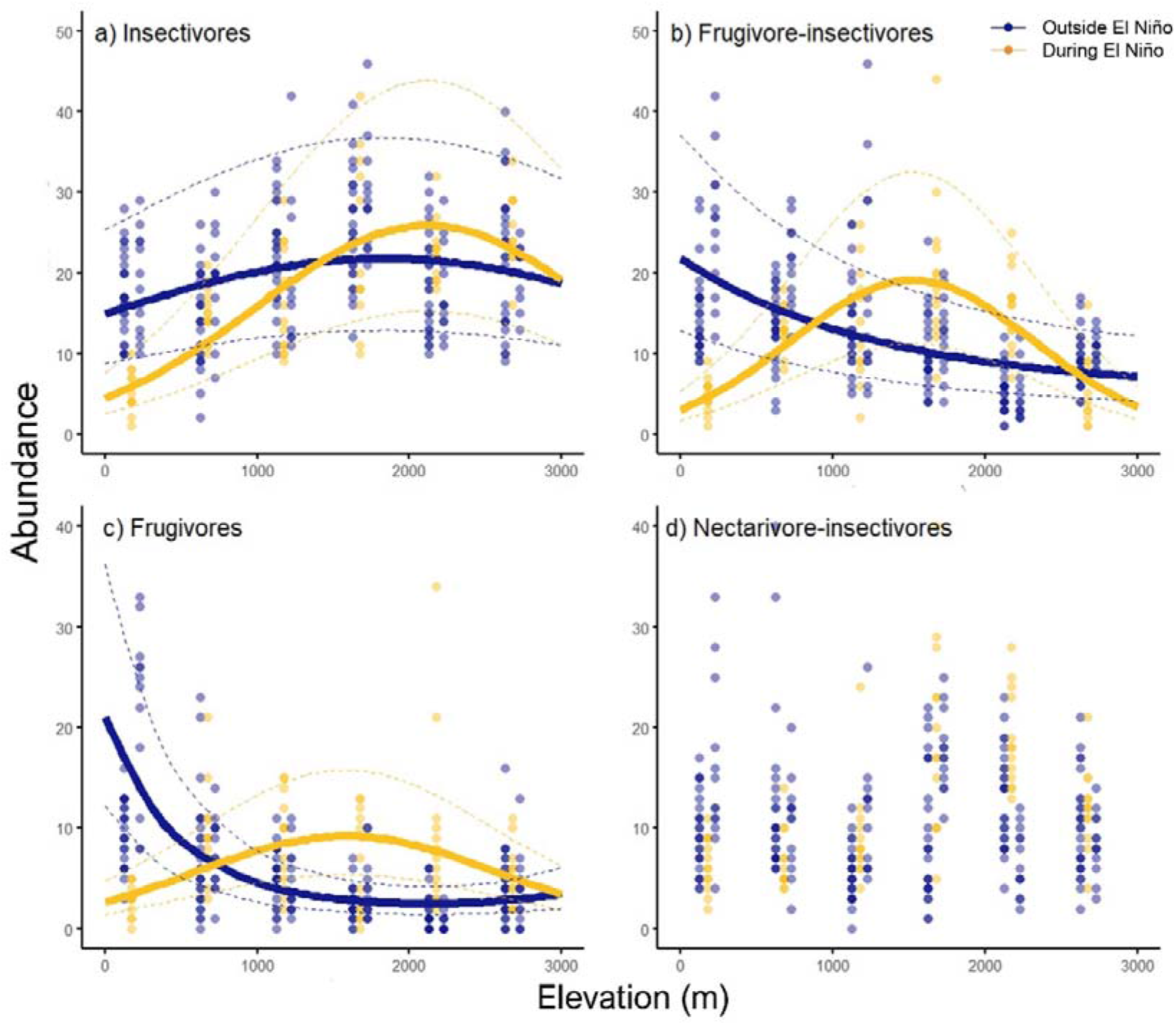
Abundance of birds belonging to individual feeding guilds recorded each time period (colour coded) at 13 points spatially distributed at each of the elevational study sites. Abundances are expressed as number of individuals recorded in 45 minutes per 0.78 ha, i.e. pooled from 3 resurveys per point with radius 50 m. Each of the points was surveyed for 15 minutes per day and the survey at each point was replicated three times in three consecutive days every year. Fitted lines represent the fit from the best model from Table 2 and dashed lines represent 95% confidence intervals of the predicted models.

**Figure 5.**
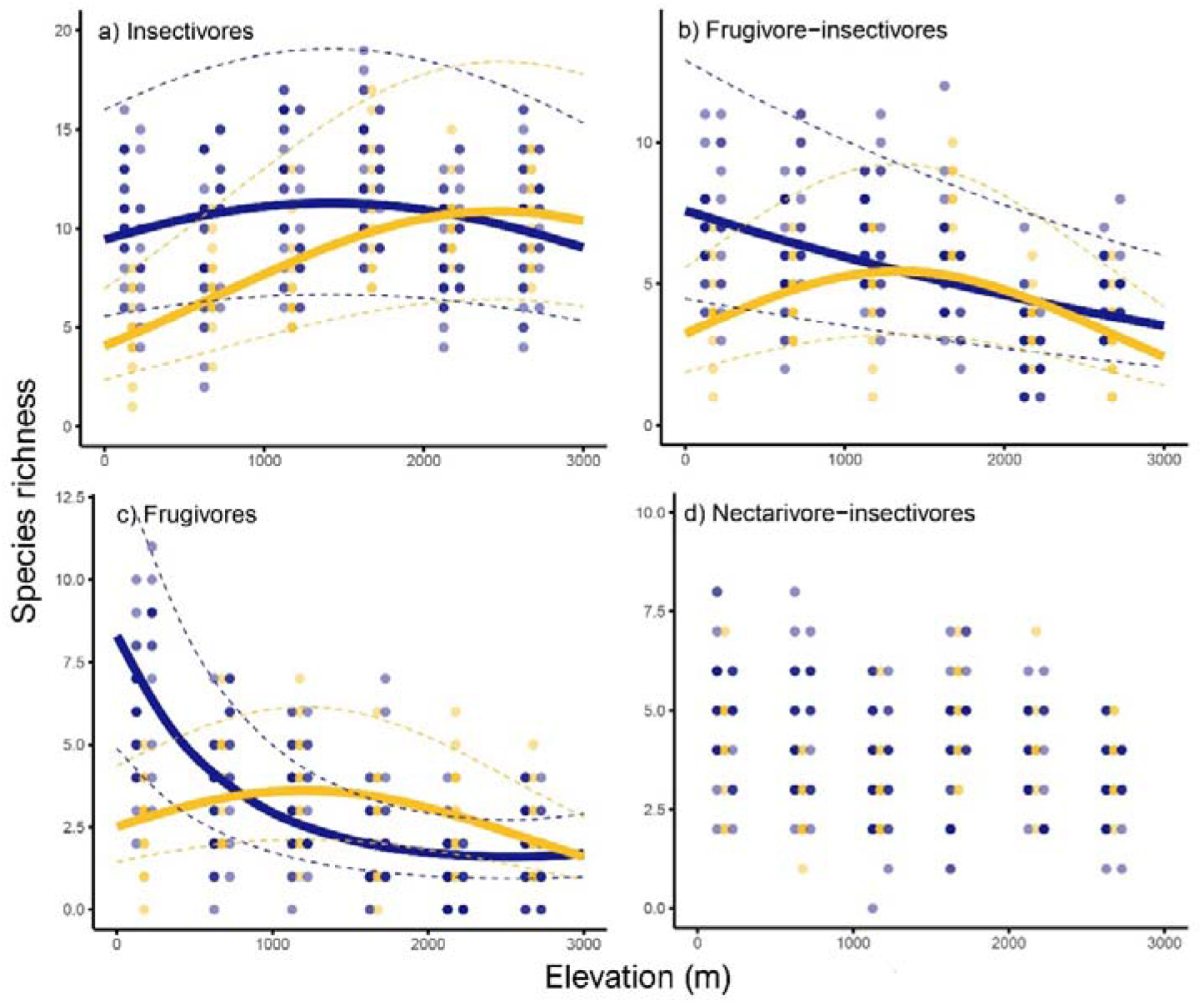
Species richness of birds belonging to individual feeding guilds recorded each time period (color coded) at 13 points spatially distributed at each of the elevational study sites. Species richness are expressed as number of individuals recorded in 45 minutes per 0.78 ha, i.e. 3 pooled surveys per point with radius 50 m. Each of the points was surveyed for 15 minutes per day and the survey at each point was replicated three times in three consecutive days every year. Fitted lines represent the fit from the best model from Table 1 and dashed lines represent 95% confidence intervals of the predicted models.

**Table 2.** Candidate models affecting abundance and richness of birds belonging to individual feeding guilds along an elevational gradient in Papua New Guinea. All models are generalized linear mixed models and include all the year/elevation combinations as a random intercept factor in addition to the fixed factors listed on the table. The null model includes only a fixed intercept in addition to the random factor. Factors and their combinations having effect on observed species richness and abundance. Effect of elevation is considered as polynomial or linear. *Long-term* effect of El Niño in models is represented by three levels – prior to, during, and after El Niño, while *resilience* is represented by 2 levels only – outside and during El Niño.

|  | Abundance |  |  | Species richness |  |  |
| --- | --- | --- | --- | --- | --- | --- |
|  | dAICc | df | weights | dAICc | df | weights |
| **Frugivores** |  |  |  |  |  |  |
| elevation<sup>2</sup> * resilience | <u>0</u> | <u>7</u> | <u>0.8516</u> | <u>0</u> | <u>7</u> | <u>0.4089</u> |
| elevation<sup>2</sup> * long-term | 3.6 | 10 | 0.1407 | 0.3 | 10 | 0.3519 |
| elevation * resilience | 6.2 | 5 | <0.0001 | 3 | 5 | 0.0912 |
| elevation * long-term | 9 | 7 | <0.0001 | 3.6 | 7 | 0.0676 |
| elevation<sup>2</sup> + resilience | 11.3 | 5 | 0.0029 | 7.2 | 5 | 0.0111 |
| elevation<sup>2</sup> + long-term | 13.3 | 6 | 0.0011 | 9.1 | 6 | 0.0043 |
| elevation + resilience | 10.8 | 4 | <0.0001 | 6.5 | 4 | 0.0089 |
| elevation + long-term | 12.8 | 5 | <0.0001 | 8.5 | 5 | 0.0058 |
| elevation<sup>2</sup> | 11.2 | 4 | 0.0031 | 5.7 | 4 | 0.0283 |
| elevation | 10.5 | 3 | <0.0001 | 4.9 | 3 | 0.0353 |
| resilience | 17.4 | 3 | <0.0001 | 21.7 | 3 | <0.0001 |
| long-term | 19.4 | 4 | <0.0001 | 23.7 | 4 | <0.0001 |
| null | 16.6 | 2 | <0.0001 | 19.9 | 2 | <0.0001 |
| **Frugivore-insectivores** |  |  |  |  |  |  |
| elevation<sup>2</sup> * resilience | <u>0</u> | <u>7</u> | <u>0.5164</u> | 0.50 | 7.00 | 0.1295 |
| elevation<sup>2</sup> * long-term | 0.20 | 10.00 | 0.4673 | 5.90 | 10.00 | 0.0087 |
| elevation * resilience | 9.40 | 5.00 | 0.0047 | 0.50 | 5.00 | 0.1295 |
| elevation * long-term | 10.40 | 7.00 | 0.0028 | 3.70 | 7.00 | 0.0261 |
| elevation<sup>2</sup> + resilience | 13.30 | 5.00 | 0.0007 | 1.30 | 5.00 | 0.0868 |
| elevation<sup>2</sup> + long-term | 13.60 | 6.00 | 0.0009 | 3.30 | 6.00 | 0.0319 |
| elevation + resilience | 12.30 | 4.00 | 0.0011 | <u>0</u> | <u>4</u> | <u>0.1662</u> |
| elevation + long-term | 12.60 | 5.00 | 0.0006 | 2.10 | 5.00 | 0.0582 |
| elevation<sup>2</sup> | 11.30 | 4.00 | 0.0018 | 1.70 | 4.00 | 0.0710 |
| elevation | 10.30 | 3.00 | 0.0030 | 0.30 | 3.00 | 0.1431 |
| resilience | 16.40 | 3.00 | 0.0001 | 8.30 | 3.00 | 0.0026 |
| long-term | 17.10 | 4.00 | <0.0001 | 10.30 | 4.00 | 0.0014 |
| null | 14.40 | 2.00 | 0.0004 | 7.80 | 2.00 | 0.0034 |
| **Insectivores** |  |  |  |  |  |  |
| elevation<sup>2</sup> * resilience | <u>0</u> | <u>7</u> | <u>0.8628</u> | 0.5 | 7 | 0.3623 |
| elevation<sup>2</sup> * long-term | 5.5 | 10 | 0.0552 | 6.4 | 10 | 0.0190 |
| elevation * resilience | 5.5 | 5 | 0.0552 | <u>0</u> | <u>5</u> | <u>0.4652</u> |
| elevation * long-term | 9.1 | 7 | 0.0091 | 3.8 | 7 | 0.0696 |
| elevation<sup>2</sup> + resilience | 10.3 | 5 | 0.0050 | 6.2 | 5 | 0.0180 |
| elevation<sup>2</sup> + long-term | 12.1 | 6 | 0.0020 | 8.6 | 6 | 0.0063 |
| elevation + resilience | 12.3 | 4 | 0.0018 | 6.6 | 4 | 0.0172 |
| elevation + long-term | 14.2 | 5 | 0.0007 | 8.7 | 5 | 0.0060 |
| elevation<sup>2</sup> | 10.1 | 4 | 0.0055 | 10 | 4 | 0.0031 |
| elevation | 11.9 | 3 | 0.0022 | 9.7 | 3 | 0.0036 |
| resilience | 17.3 | 3 | 0.0002 | 6.5 | 3 | 0.0180 |
| long-term | 19.2 | 4 | 0.0001 | 8.5 | 4 | 0.0066 |
| null | 16.5 | 2 | 0.0002 | 9.1 | 2 | 0.0049 |
| **Nectarivore-insectivores** |  |  |  |  |  |  |
| elevation<sup>2</sup> * resilience | 4.30 | 7.00 | 0.0157 | 3.70 | 7 | 0.0025 |
| elevation<sup>2</sup> * long-term | 6.90 | 10.00 | 0.0495 | 8.50 | 10 | 0.0339 |
| elevation * resilience | 4.50 | 5.00 | 0.1564 | 2.60 | 5 | 0.0152 |
| elevation * long-term | 6.40 | 7.00 | 0.0777 | 6.50 | 7 | 0.0394 |
| elevation<sup>2</sup> + resilience | 4.90 | 5.00 | 0.0100 | 0.40 | 5 | 0.2505 |
| elevation<sup>2</sup> + long-term | 5.90 | 6.00 | 0.0272 | 2.30 | 6 | 0.0435 |
| elevation + resilience | 3.90 | 4.00 | 0.1564 | 1.20 | 4 | 0.1071 |
| elevation + long-term | 4.90 | 5.00 | 0.0777 | 3.10 | 5 | 0.0394 |
| elevation<sup>2</sup> | 2.80 | 4.00 | 0.0426 | <u>0</u> | <u>4</u> | <u>0.0877</u> |
| elevation | 1.90 | 3.00 | 0.1218 | 0.60 | 3 | 0.2267 |
| resilience | 2.00 | 3.00 | 0.1102 | 5.60 | 3 | 0.0458 |
| long-term | 3.00 | 4.00 | 0.0605 | 7.50 | 4 | 0.0168 |
| null | <u>0</u> | <u>2</u> | <u>0.2579</u> | 4.90 | 2 | 0.0969 |

We recorded upper elevational range limits to be shifted by more than 500m asl in 22 bird species during El Niño year, in contrast to their typical ranges. Majority of them (N = 14) were fruit feeding species (e.g., *Cacatua galerita, Ducula pinon, Epimachus fastosus* - Table S1). Some other species either shrank the ranges from both upper and lower limits or abandoned only lower part of the range (e.g., *Myzomela rosenbergii, Coracina boyeri* – Table S1, Fig. S4). The most detectable range shifts occurred in sicklebills (*Epimachus*) and fruit doves (*Ptilinopus*), and while we recorded them a site or two higher during El Niño survey, they seemed to return back to their typical ranges soon after that. Some species (e.g., *Dicaeum geelvinkianum, Probosciger aterrimus* - Fig. S4) only redistributed their abundances and were detected in higher abundances in the upper part of its range during El Niño only. Mean difference between lower elevational limit of all birds was −18 m between 2010 and 2013, while this difference was +24 m between 2013 and 2015 and −35 m between 2010 and 2016.

## Discussion

Local species richness and abundance of forest birds along the elevational gradient of Mt. Wilhelm in Papua New Guinea was strongly influenced by the ENSO event. According to our predictions, the effect of the event varied along the elevational gradient. The results are thus also in line with studies showing that more rich and abundant bird communities are found in locations rich in rains (e.g., Rompré et al. 2007) and that birds potentially migrate to less impacted areas, or survive there more likely, during ENSO events (Blake and Loiselle 2000, Beissinger 2008, Boyle et al. 2020). Our results imply that their communities were rather resilient to the changes. This might be in line with earlier findings showing that the majority of birds (44%) fell into the group with the highest potential for recovery after environmental perturbation in tropical Australia (Isaac et al. 2009). Especially in species richness, we observed a change in the elevational pattern only during droughts related to El Niño. As soon as the rains started to appear again few months later (December 2015, pers.obs., Hirons et al. 2020, Beauchamp et al., unpubl. data), birds at the lowest elevations of the gradient had already recovered their typical species richness. Changes in abundances were however detectable for a longer time.

This increase in species richness and abundance at low elevations coincided with mass new leaf and flower flushing which was very apparent at low elevations (< 1,200 m a.s.l.). Similar observations were done elsewhere (Leighton 1983, Blake and Loiselle 2000). The mean abundances of individual bird species at the lowest elevational study site in 2016 were similar (ca. 104%) to normal conditions prior to El Niño. However, the after El Niño abundance remained significantly lower than before El Niño along the majority of the gradient.

Contrary to the low elevations, higher mid-elevations (1,700 – 2,200 m) hosted higher abundances of birds during the El Niño and seemed to be generally more stable and less affected than other elevational sites. The observed patterns were possibly partly caused by the birds shifting their upper (and sometimes also lower) ranges above their typical ranges, but more likely by redistribution of the abundances within the typical range. As we did not track movement of the birds, we can only speculate reasons behind the observed patterns. First, birds might have been really migrating, which resulted into the redistribution of abundances within their typical ranges or migrated even outside of their typical range. Other reasons might be that rainfall impacted predation, starvation, and reproduction. As most of tropical birds are described as sedentary, if other pressure happens within their patch, they typically decrease their abundances. Finally, the observed pattern might be cause but changes in vocal behaviour of the birds due to the stress, thus change in detectability which we were not able to quantify. Studies of effect of El Niño along elevational gradients are rare, but birds showed an asynchrony and delayed laying in lowland sites in contrast to highland sites of an elevational gradient in Venezuela, while the laying was synchronized in years outside of El Niño (Beissinger 2008). Combinations of strategies among Mt. Wilhelm birds is likely. During El Niño, we observed not only significant decreases in abundance of individual bird species at the lowest elevations of the gradient (200 and 700 m a.s.l.) but we also recorded several species higher, above their typical elevational ranges. After the end of El Niño, their abundances increased significantly (in contrast to El Niño conditions) at the elevations which they were using originally. The birds which we detected higher than usually were mostly frugivores (14 out of 22 bird species, Table S2), which is in line with our predictions. Frugivorous and nectarivorous species offer conspicuous exemptions of typical tropical sedentary bird and travel locally to follow patchy resources such as fruits and flowers (Blake and Loiselle 2000, Pratt and Beehler 2015). Our results further call for more robust data from Papua New Guinea (and other less studied regions), from across seasons, as the current survey can’t provide convincing reports on potential shifts of ranges or abundances of birds.

From within insectivores, several monarchs and fantails (Table S2) also seemed to undergo elevational migrations, which is rather surprising as they are described as sedentary and nonmigratory (Pratt and Beehler 2015). However, some the populations of monarch and fantails in Australia are known to be partly migratory, with some individuals being resident and some migrating regularly (Chan 2001). Evidence for facultative or regular migrations of birds along the slopes of Mt Wilhelm is assumed but mostly undescribed (Pratt and Beehler 2015).

Our data imply that the birds along Mt. Wilhelm elevational gradient might be resilient to extreme environmental events. Some tropical bird communities seemed to be resilient, but other researchers for the resilient (Isaac et al. 2009)to be rather untypical for them (Sekercioglu et al. 2002). Specific conditions of our study sites might be the reason why this study could be considered as an important count-example. We however argue that the quick recovery of the bird communities along Mt. Wilhelm might be thanks to its continuous, and mostly undisturbed, rainforest cover spanning from lowlands to high elevations. We speculate that lowland sites which do not have a ready access to higher elevations (and that is majority of lowlands of PNG) may be much more severely affected. This is especially alarming since the effects of human land use on bird diversity are typically most severe at the base of tropical mountains (Ferger et al. 2017). In a meta-analysis, Mantyka-pringle et al. (2012) found that the negative effect of habitat loss on species diversity was usually the most severe in hot areas for several taxa including birds. Because ENSO events are getting more and more severe and less predictable, and at the same time, the disturbance of forest sites increases, our results have important implications for conservation of tropical birds.

Similarly to our observations, Jaksic and Lazo (1999), Styrsky and Brawn (2011) or review of Boyle et al. (2020) reported that birds apparently reached their peak diversity and density in wet years associated to El Niño, and abundances and richness of birds were usually positively associated with rainfalls. The authors explained these increases as a consequence of augmented productivity of vegetation and secondarily also of higher arthropod abundance. In our study, the rains after the dry El Niño resulted in an increase by ca. 70% in abundance and by ca. 45% in richness in contrast to the El Niño year at most of the affected sites. Our study further suggests that increased productivity reaches the birds in the second trophic level (i.e. consumers of plants) already within six months after El Niño-driven rain. In another study, carnivorous predators (hawks, owls) displayed a one-year delayed response to small mammal abundances in another study (Jaksic 2001).

Our measurements of temperature and humidity revealed that these variables did not change too much at higher elevations, and thus provided conditions very similar to those in other years. We could observe during the field work that while higher elevations (1700 and 2700 m) were still moist maybe thanks to the cloud layer, the lower elevations were seriously affected by droughts during our visit in October 2015. A study focusing on the impact of El Niño on the agriculture also concluded that impact of severe droughts in the highlands was somewhat mitigated by a cloud layer (ca. 1700 – 2200 m a.s.l.; Hazenbosch pers. com.). The local villagers also reported significant wilt on their gardens at study sites < 700 m a.s.l., higher abundances of pests at 1200 – 2200 m a.s.l., and no changes in crop growth (or local frosts) were reported at higher elevational study sites (Beauchamp et al., unpubl. data). In another unpublished experiment (Sam et al., unpubl. data), we observed 25% and 15% mortality of experimental seedlings at 200 m a.s.l. in 2015 which was significantly higher than 0.5-1% at high elevations (Wagia pers. com.). Unfortunately, we were not able to relate our bird data to changes in primary productivity of food resources more precisely.

El Niño/Southern Oscillation constitutes an excellent natural experiment and at the same time illustrates how atmospheric-oceanographic phenomena may affect terrestrial biota. It can be used as a tool to understand responses to climate change, and as itself a result from climate change in its intensity and frequency. The alternate phases of ENSO can be studied to make reasonable extrapolations about where global climate change may lead us. However, future studies would need to focus both on during El Niño as well as outside of Niño surveys, which would allow us to analyse more exactly what happens with bird communities after and during the disturbance. Warm ENSO events were historically very frequent over the Holocene until about 1,200 years ago, and then declined towards the present (Moy et al. 2002). However, the report showing that four out of 12 strong Niño events occurred in the last 20% of the 20th century (1982-83, 1986-87, 1991-92 and 1997-98) (Jaksic 2001), suggests that we should be more concerned about the increasingly more extreme events of El Niño, rather than about a gradual warming of the climate (Jaksic 2001). Also an occurrence might be of concern as the situation is that we have problem to predict an El Niño event few months in future. There were 16 El Niño alerts but only 10 events since 1980.

## Acknowledgement

We are thankful to villagers from Kausi, Numba, Degenumbu, Sinopass, Bruno Sawmill and Kegesugl for allowing work on their land and assistance in the field. The work was supported by Czech Science Foundation Grant 18-23794Y and European Research Council Starting Grant BABE 805189 (PI: K. Sam). We are thankful to Michael Kigl, Edwin Sohun and Penniel Lamei for help in the field.

## Notes

### Competing Interest Statement

The authors have declared no competing interest.

