## Supplement for "Reorganization of bird communities along a rainforest elevation gradient during a strong El Niño event in Papua New Guinea"

**Appendix S1** Additional methods, R codes and list of species

Figure S1 - Forest interior and canopy openness at elevational sites. Each picture represents habitat with the mean score for given elevation: 200 m – shrub density 8%, canopy openness 11%; 700 m – shrub density 12%, canopy openness 15%; 1200 m - shrub density 39%, canopy openness 16%; 1700 m - shrub density 40% (note the track); canopy openness 17%, 2200 m - shrub density 38%; canopy openness 40%; 2700 m - shrub density 34%, canopy openness 40%; 3200 m - shrub density 38%, canopy openness 21%; 3700 m - shrub density 20%, canopy openness 90%. Pictures are only illustrative as the measurements for each variable were averaged for each point.

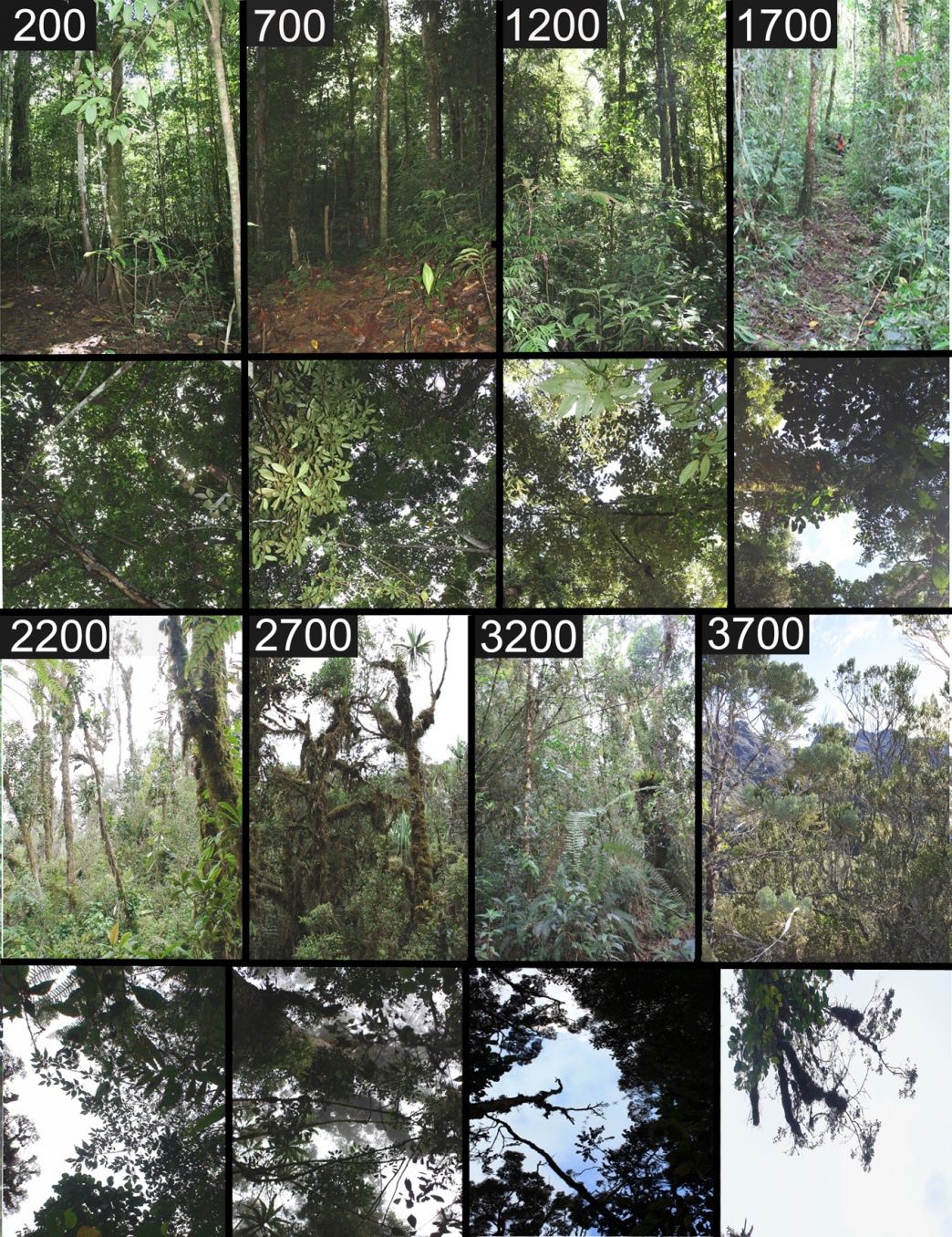

**Figure S2**. Correlation between averaged, summed, and maximal abundances across the three-consecutive point-counts.

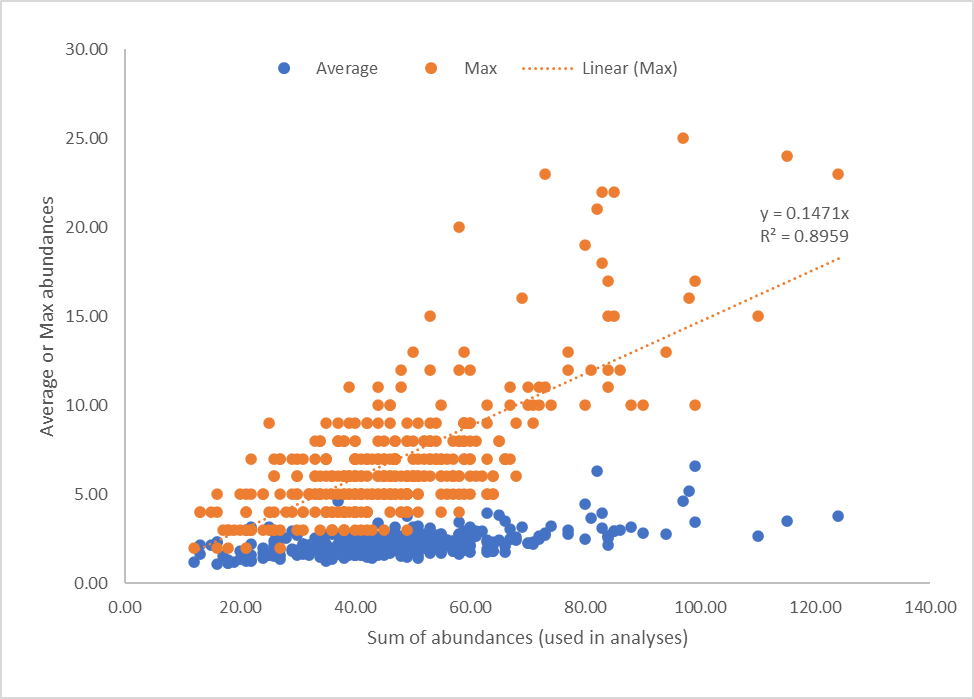

**Figure S3.** Mean Chao 1 estimates of species richness, its lower and upper 95% confidence intervals and raw observed number of species.

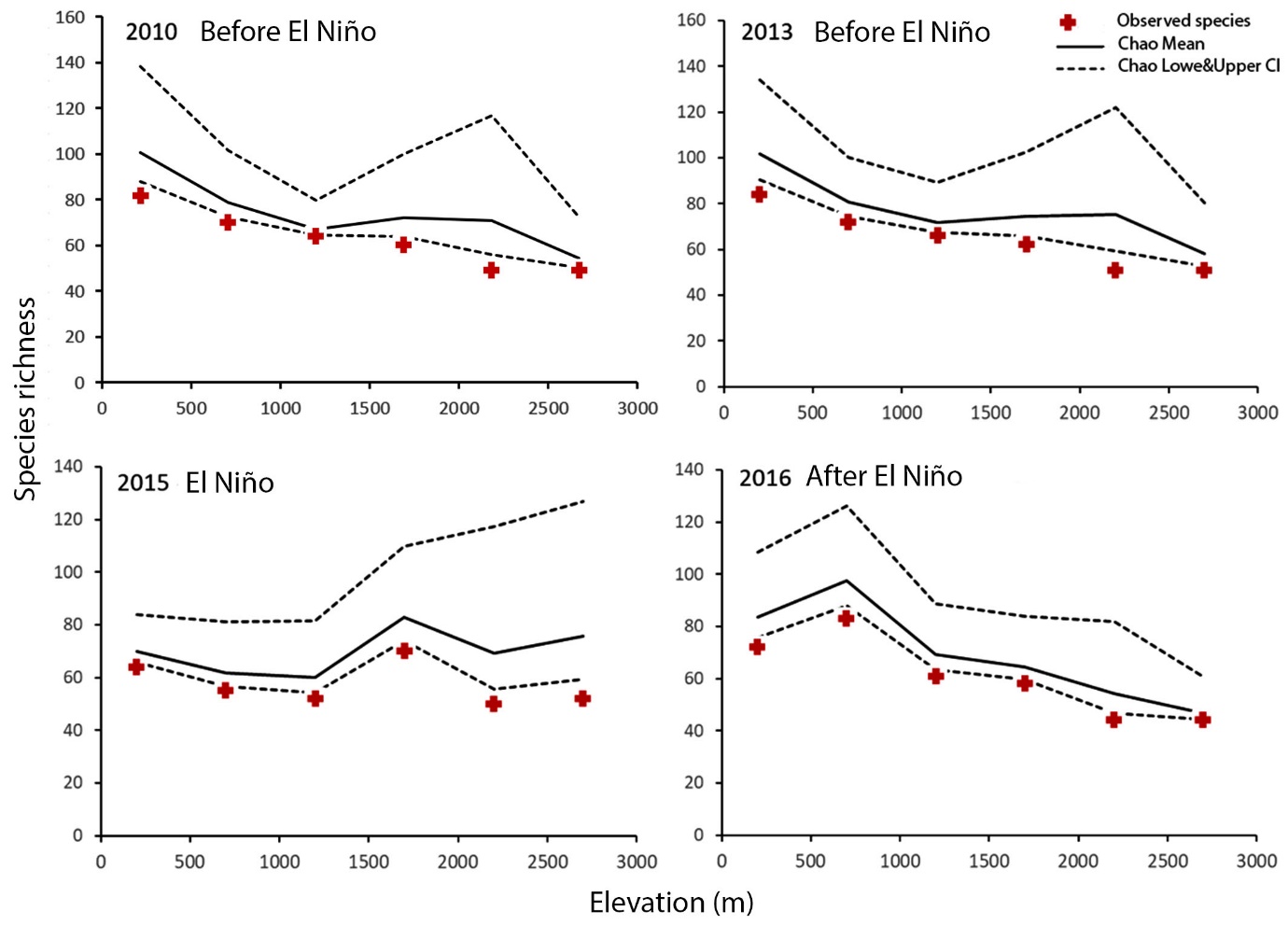

**Table S1** – List of recorded bird species and their ocurences at studied sites in individual years. Typical ranges were extracted from Hanbook of the Birds of the World (online version) and are marked in green. Black rectangles mark the presence of bird on given elevation each survey. Birds marked by * extended their distribution beyond their typical range in El Nino year.

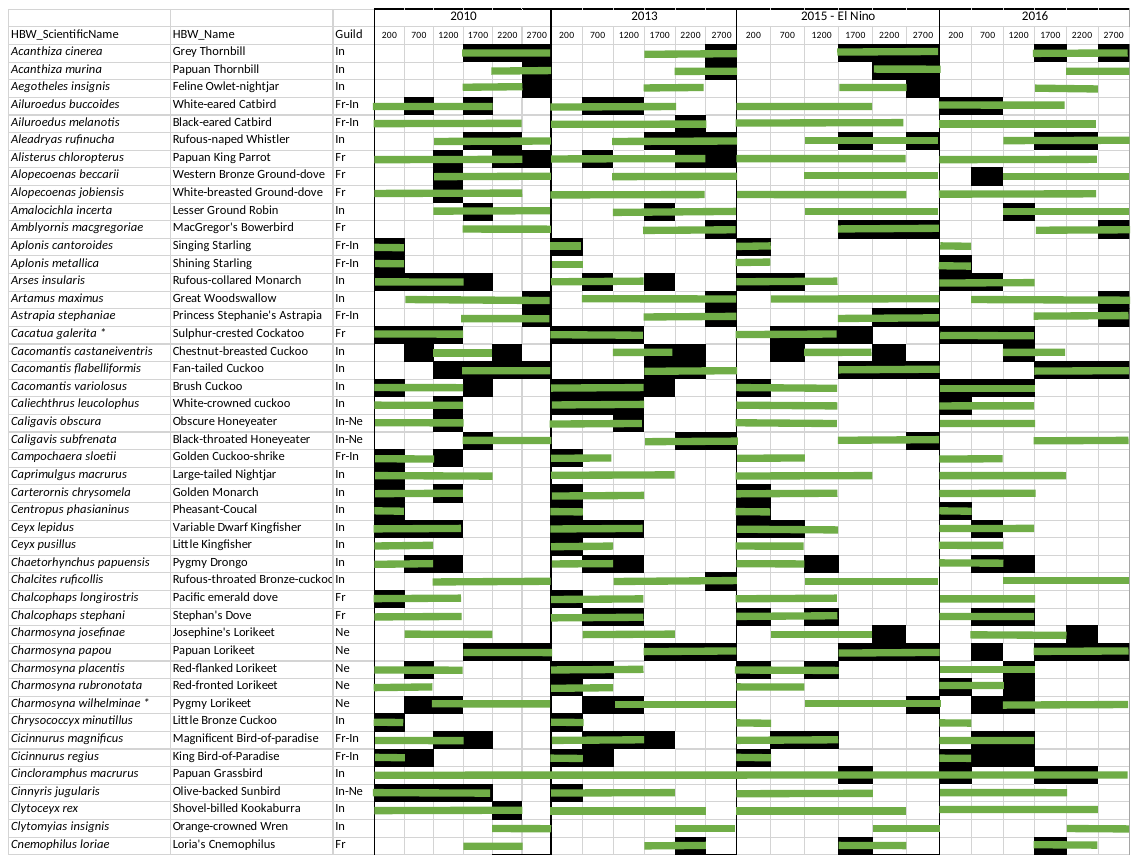

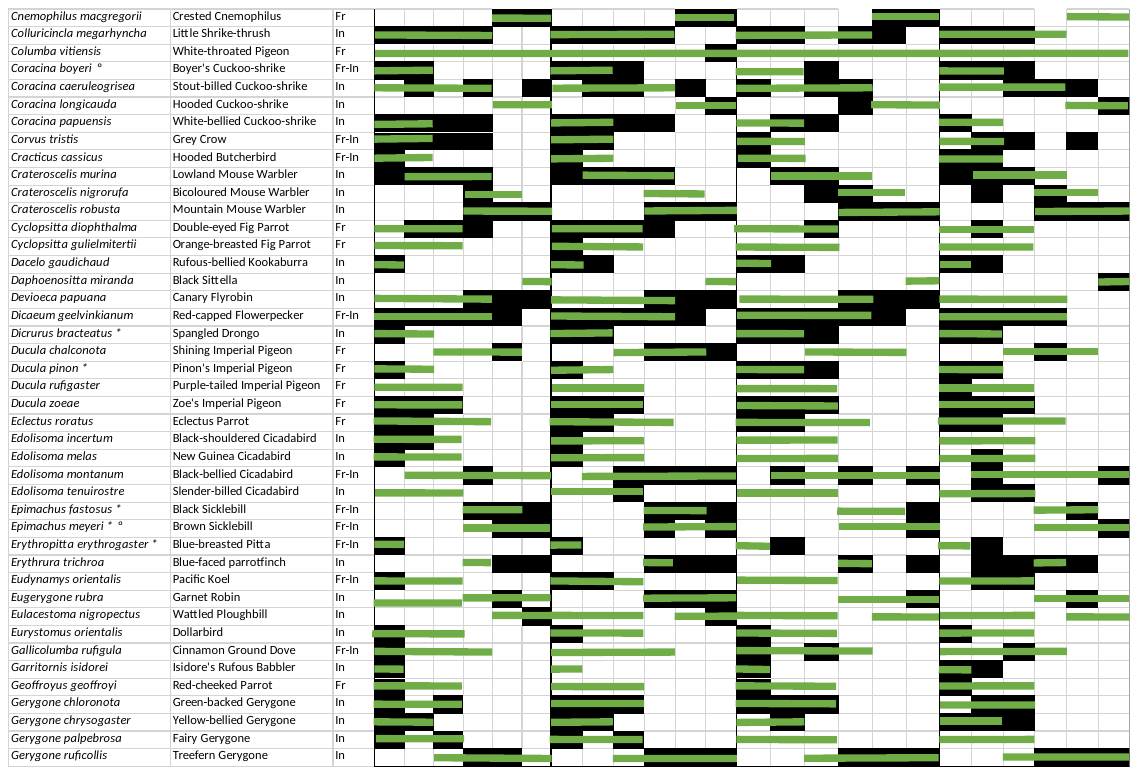

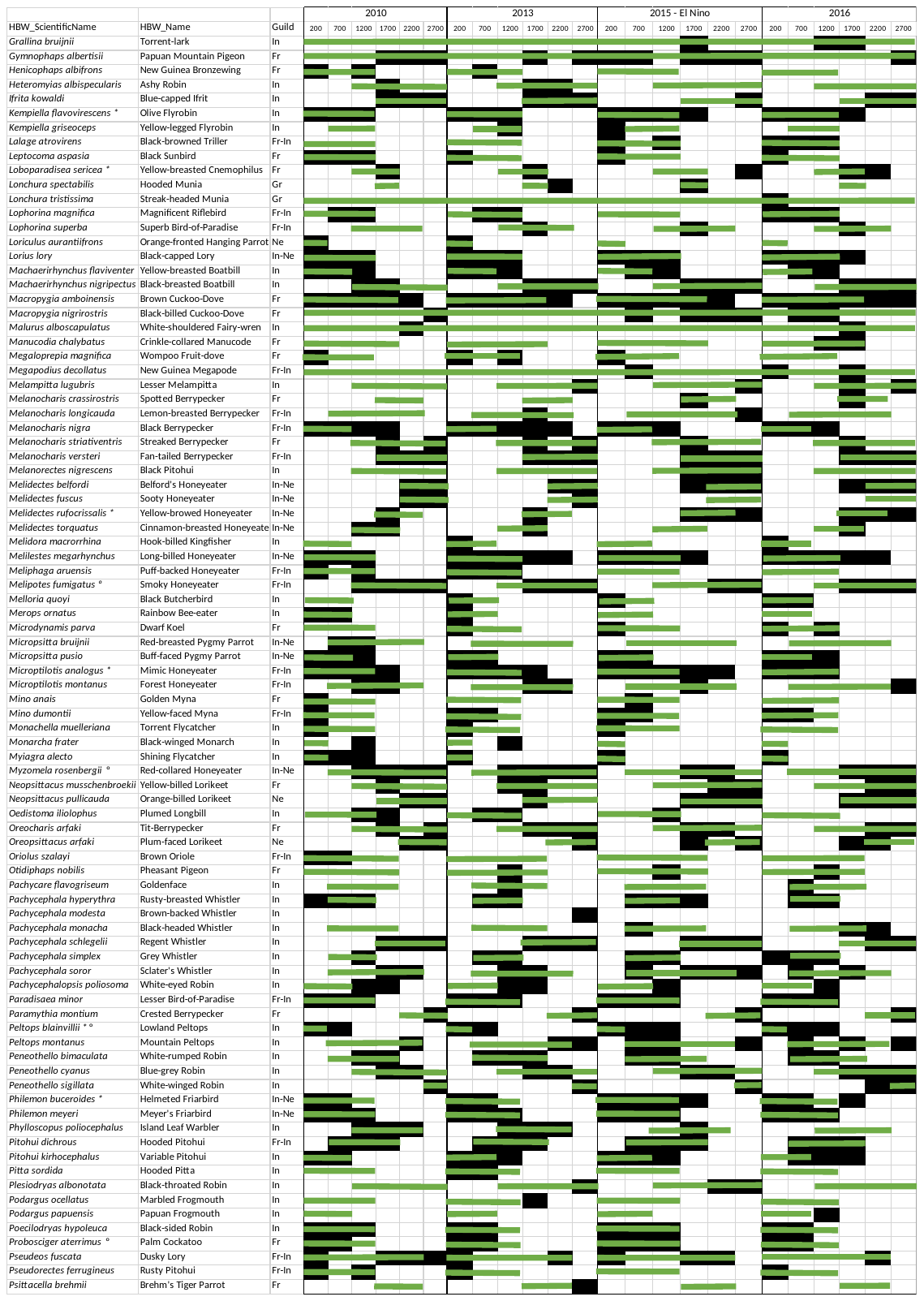

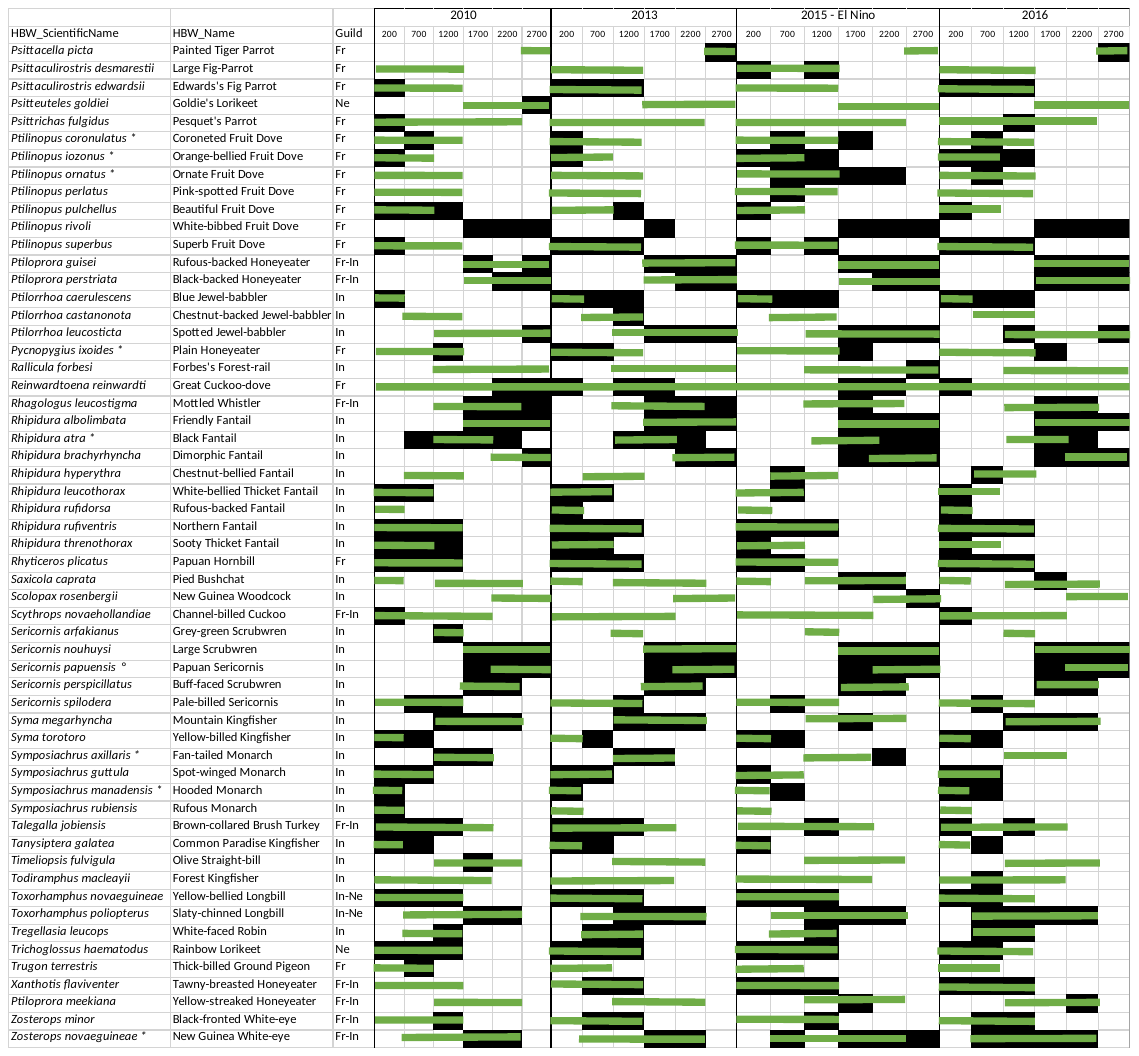

Birs of prey and swifts excluded from analyses: *Accipiter melanochlamys*, *Accipiter meyerianus*, *Elanus caeruleus*, *Haliastur indus*, *Haliastur sphenurus*, *Harpyopsis novaeguineae*, *Henicopernis longicauda*, *Milvus migrans*, *Accipiter novaehollandiae, Apus pacificus*

**Table S2.** Summary and confidence intervals of the effects from the full models.

|  | Abundance | | | | | |
| --- | --- | --- | --- | --- | --- | --- |
|  | Estimate | Std. Error | z value | P | Lower CI | Upper CI |
| elevation | -0.999 | 0.506 | -1.975 | 0.048 | -2.036 | 0.019 |
| elevation^2^ | 0.349 | 0.505 | 0.691 | 0.490 | -0.683 | 1.381 |
| resilience | -0.002 | 0.050 | -0.035 | 0.972 | -0.104 | 0.100 |
| long-term | -0.149 | 0.050 | -2.972 | 0.003 | -0.252 | -0.047 |
| elevation^2^ : resilience | 6.197 | 0.897 | 6.911 | <0.001 | 4.373 | 8.031 |
| elevation : resilience | -5.333 | 0.890 | -5.992 | <0.001 | -7.154 | -3.520 |
| elevation : long-term | -1.848 | 0.885 | -2.089 | 0.037 | -3.660 | -0.045 |
| elevation^2^ : long-term | 0.225 | 0.882 | 0.255 | 0.799 | -1.574 | 2.033 |
|  | Species richness | | | | | |
| elevation | -2.316 | 0.624 | -3.709 | <0.001 | -3.563 | -1.457 |
| elevation^2^ | 0.042 | 0.620 | 0.068 | 0.946 | -0.954 | 1.142 |
| resilience | -0.147 | 0.062 | -2.367 | 0.018 | -0.276 | -0.035 |
| long-term | 0.028 | 0.061 | 0.463 | 0.643 | -0.105 | 0.162 |
| elevation^2^ : resilience | -1.695 | 1.094 | -1.549 | 0.121 | -3.874 | 0.391 |
| elevation : resilience | 4.107 | 1.101 | 3.729 | <0.001 | 2.156 | 6.442 |
| elevation : long-term | -0.568 | 1.079 | -0.526 | 0.599 | -2.936 | 1.789 |
| elevation^2^ : long-term | 0.146 | 1.074 | 0.136 | 0.892 | -2.531 | 0.417 |

**Figure S4.** Changes in abundances of selected bird species that seemed to redistribute their abundances within their typical elevational range.

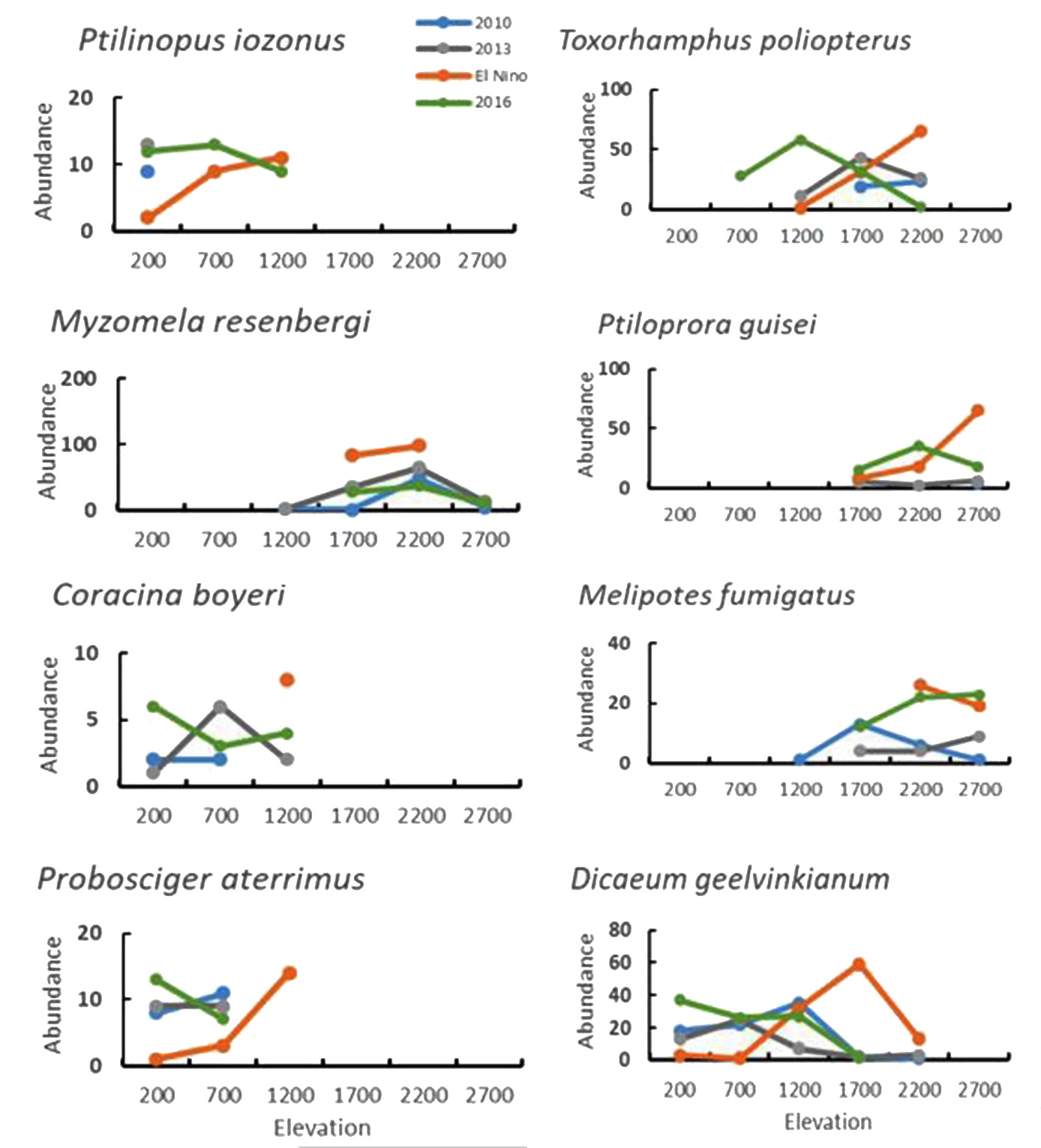
